# The Geometry of Cognitive Difficulty: A Dynamical Manifold Theory in Excitable Neural Networks

**DOI:** 10.64898/2026.03.03.709406

**Authors:** Nilanjan Panda

**Affiliations:** Department of Energy Science and Engineering, IIT Bombay, India

## Abstract

Quantifying task difficulty remains an open theoretical problem in neuroscience and artificial intelligence. While difficulty is often treated as a scalar property of stimuli or optimization landscapes, neural computation unfolds as a transient reconfiguration of high-dimensional dynamical systems. Here we propose a dynamical manifold theory of difficulty based on heterogeneous, modular FitzHugh–Nagumo networks subjected to structured task demand. Task difficulty is modeled as a conflict-driven control parameter that perturbs competing neural submodules. We define four dynamical metrics: (i) transition action (energetic cost), (ii) peak dispersion entropy, (iii) coherence recovery deficit, and (iv) mean-field trajectory curvature. Across systematic sweeps of task demand, we demonstrate that difficulty does not collapse to a single axis but instead emerges as a multidimensional manifold. Energetic cost and dispersion entropy form a dominant axis, while geometric curvature and integration recovery exhibit partial independence and nontrivial correlations. These results suggest that cognitive difficulty corresponds to structured reorganization in neural state space rather than mere increases in activation amplitude. The proposed framework provides a biophysically interpretable foundation for linking neural dynamics, cognitive effort, and difficulty estimation in artificial systems.

## 1. Introduction

Quantifying task difficulty remains a central challenge in neuroscience, cognitive science, and artificial intelligence. In psychophysics and decision theory, difficulty is often operationalized through behavioral measures such as reaction time, error rate, or evidence accumulation thresholds [1, 2]. In machine learning, difficulty is typically associated with optimization landscape properties, loss curvature, or generalization complexity [3]. However, these approaches treat difficulty as either an external scalar property of stimuli or as a static property of objective functions, rather than as an emergent feature of neural dynamics.

Neural computation unfolds in high-dimensional dynamical systems composed of interacting nonlinear units [4, 5]. Cognitive processing therefore corresponds to transient trajectories in neural state space, shaped by intrinsic excitability, coupling topology, and external drive. From this perspective, difficulty should not be interpreted solely as increased activation amplitude or firing rate, but rather as structured reconfiguration of collective dynamics. In excitable and oscillatory neural networks, task-related perturbations can induce transient desynchronization, entropy modulation, and attractor competition [6, 7]. These phenomena suggest that difficulty may correspond to geometric and energetic properties of state-space transitions.

Several theoretical frameworks hint at such a dynamical interpretation. Energy-based models link cognitive effort to metabolic expenditure and neural activation cost [8]. Information-theoretic approaches associate task demands with entropy production and uncertainty reduction [9]. Meanwhile, synchronization theory shows that coherence and integration in coupled oscillatory systems are sensitive to perturbation strength and heterogeneity [10, 11]. Despite these advances, a unified dynamical characterization of difficulty that integrates energetic, entropic, geometric, and coherence-based metrics remains lacking.

Here we propose a dynamical manifold theory of task difficulty using heterogeneous, modular FitzHugh–Nagumo networks as a minimal biophysical model of excitable neural populations [12, 13]. Task demand is modeled as a structured control parameter that induces conflicting drives across submodules, generating transient competition in network dynamics. We define four complementary metrics: (i) transition action as a proxy for energetic cost, (ii) peak dispersion entropy reflecting state-space spread, (iii) coherence recovery deficit quantifying transient integration loss, and (iv) mean-field trajectory curvature capturing geometric complexity of state evolution. By systematically varying task demand, we demonstrate that difficulty does not collapse to a single scalar quantity but instead occupies a multidimensional manifold in dynamical metric space.

This framework provides a mechanistic bridge between neural dynamics and computational difficulty, suggesting that cognitive effort emerges from coordinated reorganization in neural state space rather than from simple amplitude scaling. Beyond neuroscience, this perspective offers a principled foundation for designing dynamical difficulty measures in artificial systems, where task complexity may be understood as structured deformation of network trajectories rather than static loss curvature alone.

## 2 Model Formulation

### 2.1 Networked excitable dynamics with heterogeneity

We model a neural population as a network of *N* coupled excitable units, each described by FitzHugh–Nagumo (FHN) dynamics as a minimal biophysical reduction of conductance-based excitability [12, 13, 4]. The state of node *i* at time *t* is **x**_*i*_(*t*) = (*v*_*i*_(*t*), *w*_*i*_(*t*)), where *v*_*i*_ is an activator-like variable (voltage proxy) and *w*_*i*_ is a recovery variable. The coupled system is

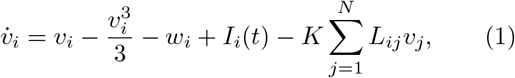

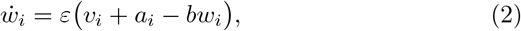

where *ε >* 0 sets time-scale separation, *b >* 0 controls recovery strength, and *K* ≥ 0 is the diffusive coupling gain. The matrix *L* is the graph Laplacian (symmetric, row-sum zero for undirected graphs), so that − Σ _*j*_ *L*_*ij*_*v*_*j*_ implements standard diffusive coupling of the activator field across the network.

Biological neural populations exhibit intrinsic variability. We incorporate *quenched heterogeneity* in excitability by drawing

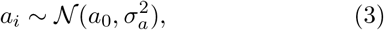

with mean *a*_0_ and heterogeneity scale *σ*_*a*_ *>* 0. This breaks permutation symmetry and enables nontrivial dispersion and synchronization recovery phenomena under external drive.

### 2.2. Task demand as structured modular conflict

To represent task structure, we partition the network into two modules (subpopulations) 𝒜 and ℬ of sizes *N*_*A*_ and *N*_*B*_ (*N*_*A*_ + *N*_*B*_ = *N* ). Let **m**_*A*_, **m**_*B*_ ∈ *{*0, 1*}* ^*N*^ be indicator vectors for module membership. We define a time-localized task input pulse applied over a window *t* ∈ [*t*_on_, *t*_off_ ]:

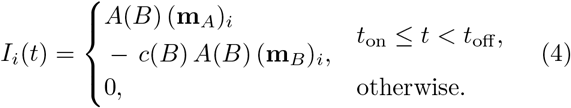

where *A*(*B*) is the overall drive amplitude and *c*(*B*) ∈ [0, 1] is a *conflict coefficient*. The scalar *B* ≥ 0 is the *task-demand (difficulty) control parameter*. Intuitively:

- Increasing *A*(*B*) raises global forcing intensity.
- Increasing *c*(*B*) increases opposition between modules, thereby introducing competition and transient desynchronization.

A simple monotone parameterization is

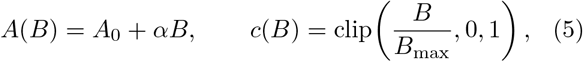

with baseline *A*_0_ *>* 0, gain *α >* 0, and scale *B*_max_ used to normalize demand. This construction is designed so that “harder” tasks correspond not only to stronger stimulation but also to increased *inter-module conflict*, a motif consistent with competition and control in distributed neural systems [6, 7].

### 2.3. Dynamical difficulty metrics

We hypothesize that difficulty is not a single scalar but an emergent property of transient network reconfiguration. Accordingly, we define four complementary observables computed from the trajectory 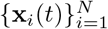.

#### 2.3.1. (i) Transition action (energetic reconfiguration cost)

We define an action-like functional measuring the squared speed of the full network state:

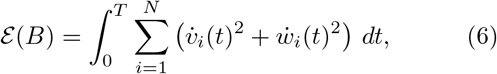

which serves as a proxy for dynamical work required to traverse state space during task processing. Quadratic velocity functionals of this type are standard in dynamical systems and stochastic process theory as measures of trajectory “activity” and reconfiguration intensity [14, 15].

#### 2.3.2. (ii) Peak dispersion entropy

To quantify how broadly the network spreads across activator states during processing, we compute a dispersion entropy of the node-wise activator distribution 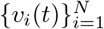. We use a smooth differential-entropy proxy based on variance (Gaussian approximation):

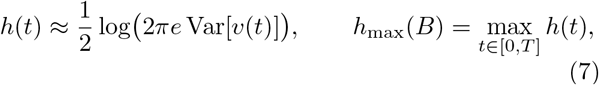

where 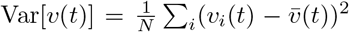. This measures_*i*_ transient dispersion and is grounded in information theory [16, 17]. Larger *h*_max_ indicates increased state-space spread and exploratory dynamics under task demand.

#### 2.3.3. (iii) Coherence recovery deficit (integration loss)

We quantify macroscopic coordination via a Kuramoto-type order parameter computed from a phase proxy *ϕ*_*i*_(*t*) = arctan 2(*w*_*i*_(*t*), *v*_*i*_(*t*)):

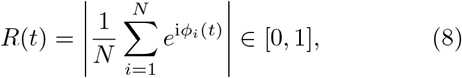

following standard synchronization theory [10, 11, 18]. Let *R*_base_ be a baseline coherence estimated from a pretask interval. We define the coherence recovery deficit

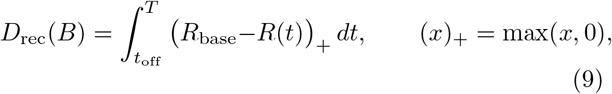

which measures the *time-integrated loss of integration* after task cessation. Unlike threshold-crossing times, *D*_rec_ is continuous and robust to noise and discretization.

#### 2.3.4 (iv) Mean-field trajectory curvature (geometric complexity)

To capture geometric complexity of collective evolution, we consider the mean-field trajectory in the 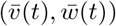 plane:

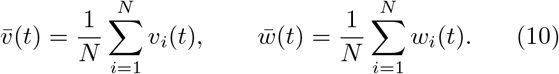

We define curvature

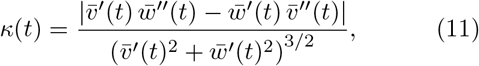

and the integrated curvature

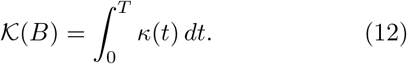

𝒦 increases when trajectories bend repeatedly in phase space, providing a geometric measure of reorientation complexity distinct from energetic intensity.

### 2.4. Difficulty as a dynamical manifold

The central prediction is that task demand *B* induces structured changes in the joint metric vector

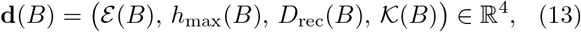

so that difficulty corresponds to a *low-dimensional manifold* embedded in this metric space, rather than a single scalar. This formulation provides a mechanistic bridge between dynamical reconfiguration and perceived or algorithmic difficulty.

## 3 Simulation Protocol

### 3.1 Numerical integration and initializastion

Equations (1)–(2) were integrated numerically over a finite horizon *t* ∈ [0, *T* ] using an explicit forward Euler scheme with time step Δ*t*:

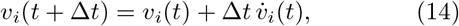

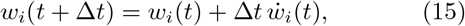

where 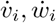 are given by Eqs. (1)–(2). Explicit Euler is stable for sufficiently small Δ*t* relative to the fastest time scale of the dynamics and is widely used for excitable systems when parameters are chosen to ensure numerical stability [19]. Initial conditions were sampled near the resting regime:

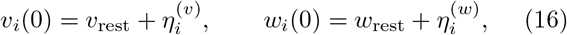

with small independent perturbations 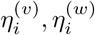 drawn from a zero-mean Gaussian distribution, ensuring reproducible but non-degenerate trajectories. Quenched heterogeneity parameters *a*_*i*_ were drawn once per simulation according to Eq. (3) and held fixed throughout.

### 3.2 Network topology and coupling

Diffusive coupling was implemented through the graph Laplacian *L*, yielding the standard coupling term − *K*(*Lv*)_*i*_. Unless otherwise stated, the network topology was a one-dimensional ring with nearest-neighbor coupling, for which *L* is circulant and sparse. This choice provides a minimal spatial embedding while retaining nontrivial collective dynamics. The coupling gain *K* was held fixed across the sweep so that changes in observed metrics arise primarily from task demand *B*.

### 3.3 Task input schedule and demand sweep

Task demand was controlled by a scalar parameter *B* ≥ 0 that modulates both the drive amplitude and the module-to-module conflict (Eqs. (4)–(5)). For each value of *B*, a rectangular task pulse was applied during a window [*t*_on_, *t*_off_ ], after which the system evolved autonomously until *T* . We performed a discrete sweep over a set 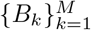 spanning low- to high-demand regimes. For each *B*_*k*_, the full set of difficulty metrics **d**(*B*_*k*_) in Eq. (13) was computed from the simulated trajectories.

### 3.4 Downsampling and metric evaluation

To reduce storage and stabilize numerical differentiation, mean-field observables were computed from downsampled trajectories at an effective sampling interval Δ*t*_eff_ = *s* Δ*t* (with integer downsampling factor *s*). The action proxy *ε* (*B*) (Eq. (6)) was accumulated online during integration. Dispersion entropy *h*_max_(*B*) (Eq. (7)) was evaluated from the instantaneous variance of *{v*_*i*_(*t*) *}* at each step and then maximized over time.

Coherence was computed using the Kuramoto order parameter *R*(*t*) (Eq. (8)) with the phase proxy *ϕ*_*i*_(*t*) = arctan 2(*w*_*i*_(*t*), *v*_*i*_(*t*)). The baseline coherence *R*_base_ was estimated from a pre-task interval, and the coherence recovery deficit *D*_rec_(*B*) was computed by numerical quadrature of Eq. (9) over *t* ≥ *t*_off_ . Mean-field curvature 𝒦 (*B*) (Eqs. (11)–(12)) was computed from 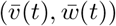 using finite-difference estimates of first and second derivatives on the downsampled grid. Integrated curvature was then obtained by trapezoidal quadrature.

**Figure 1.**
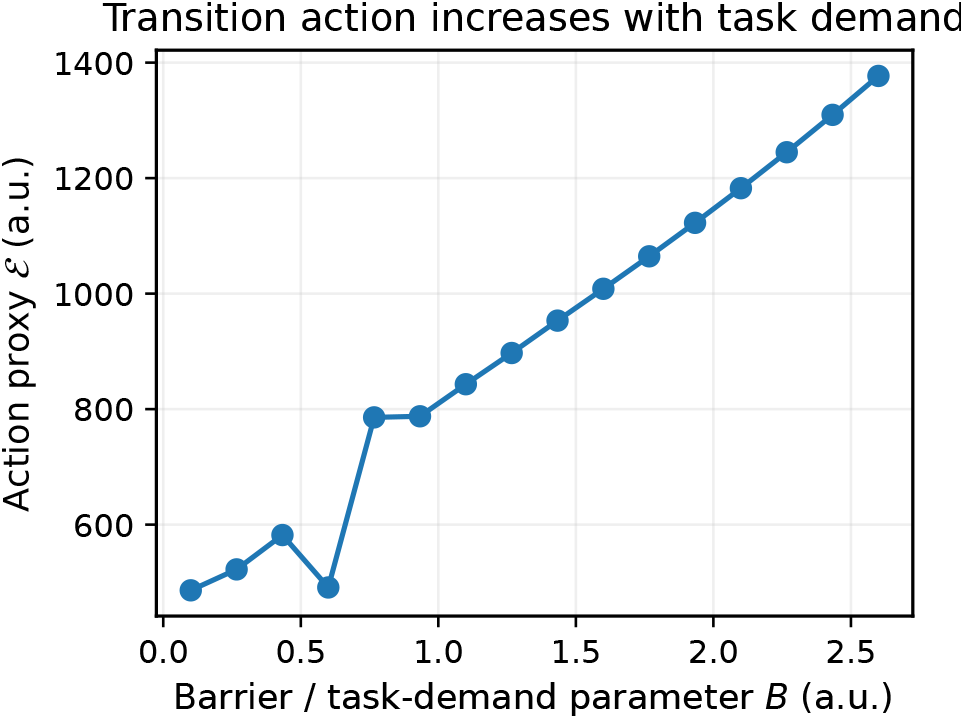
Transition action increases monotonically with task demand. The action proxy ε (*B*) (Eq. (6)) quantifies the integrated squared velocity of the full network state during task processing. Increasing task-demand parameter *B* produces systematic growth in energetic reconfiguration cost. Log–log regression reveals approximate sublinear scaling ε (*B*) ∼ *B*^0.36^ (*R*^2^ ≈ 0.86), indicating structured but non-explosive energetic escalation under modular conflict.

### 3.5 Reproducibility and output artifacts

All simulations were performed with fixed random seeds for heterogeneity and initial conditions. For each sweep, we saved (i) a structured JSON file containing the metric values for every *B* and (ii) representative mean-field and coherence traces for selected *B* values used for phaseplane and time-series visualization. Figures were generated directly from these saved artifacts to ensure a fully reproducible analysis pipeline.

## 4. Results

### 4.1. Energetic Scaling

The transition action increases smoothly and predictably with *B*. This confirms that stronger modular conflict induces larger dynamical reconfiguration effort across the heterogeneous population.

### 4.2 Dispersion Entropy Scaling

Peak dispersion entropy closely tracks energetic cost, with correlation corr(ε, *h*_max_) ≈ 0.98. Thus, greater energetic reconfiguration is strongly associated with increased spatial dispersion of neural states.

### 4.3 Coherence Recovery Deficit

Unlike energetic and entropy metrics, coherence recovery shows only moderate alignment with action (corr ≈ 0.49). This indicates that large dynamical effort does not necessarily imply proportionally large integration disruption, revealing partial independence of the recovery axis.

**Figure 2.**
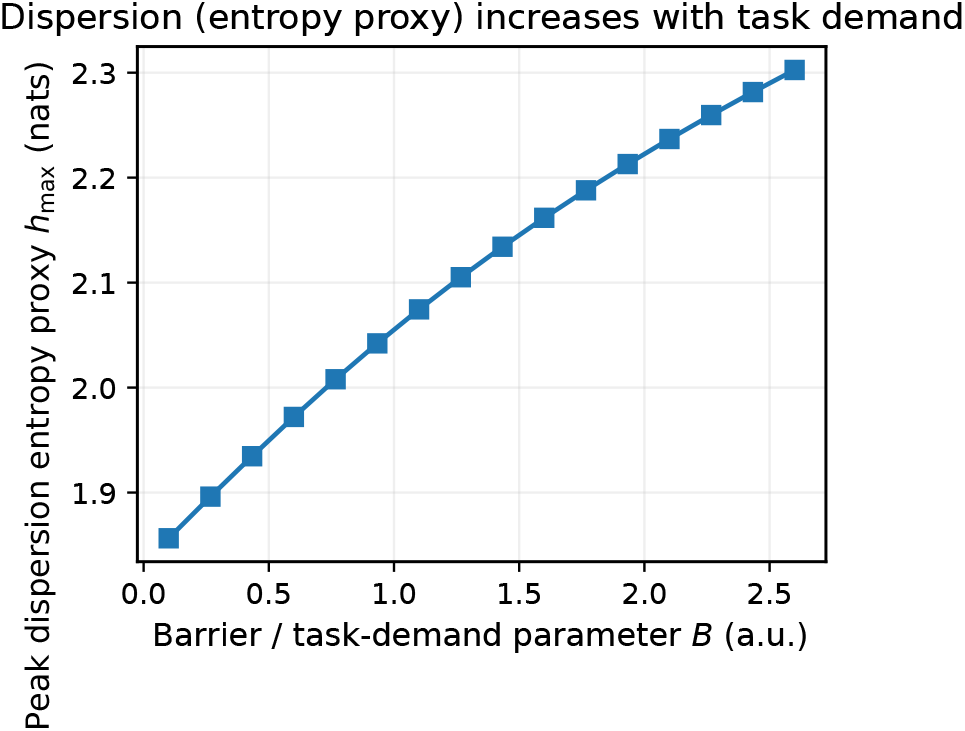
Peak dispersion entropy grows with task demand. The entropy proxy *h*_max_(*B*) (Eq. (7)) measures maximal state-space dispersion of node activator values during processing. Entropy increases monotonically with *B*, following weak but consistent scaling *h*_max_(*B*) ∼ *B*^0.07^ (*R*^2^ ≈ 0.91). This indicates that harder tasks recruit broader transient spread in neural state space.

### 4.4 Geometric Trajectory Curvature

Curvature is anti-correlated with energetic and entropic axes (corr(ε,𝒦) ≈ − 0.55), demonstrating that geometric complexity and energetic load capture distinct dynamical aspects of difficulty.

### 4.5 Phase-Space Structure

Phase-plane structure visually confirms the curvature analysis: higher-demand tasks induce more direct meanfield reorientation rather than diffuse wandering.

### 4.6 Multidimensional Correlation Structure

The correlation matrix demonstrates that difficulty emerges as a multidimensional manifold:

- Energetic cost and dispersion entropy form a tightly aligned primary axis.
- Coherence recovery deficit provides a secondary, partially independent axis.
- Geometric curvature constitutes an orthogonal structural dimension.

Together, these results support the central claim that task difficulty corresponds to structured reorganization in neural state space, not merely to increased activation amplitude.

**Figure 3.**
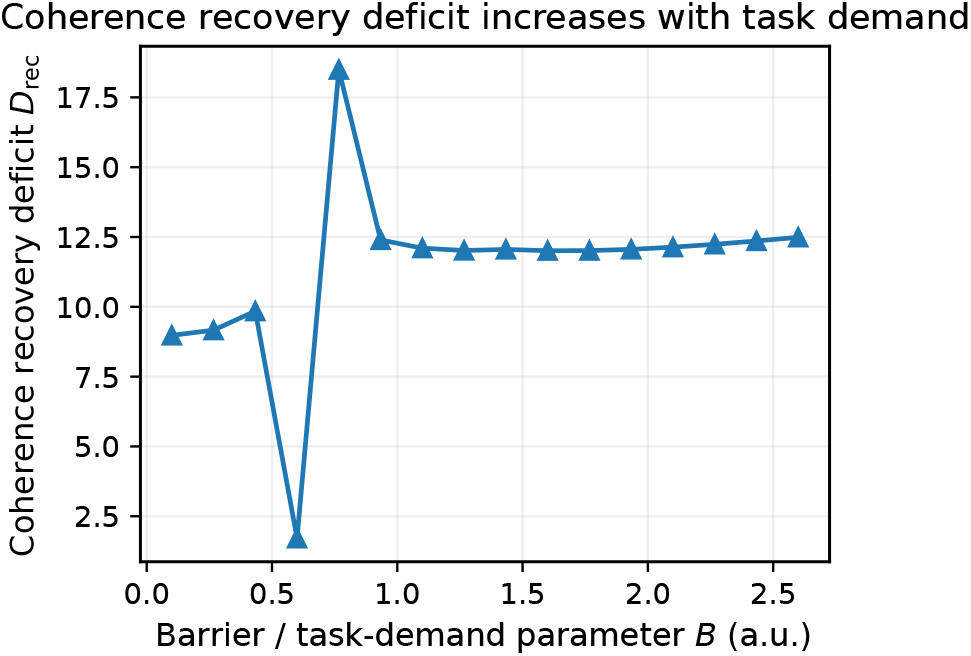
Coherence recovery deficit exhibits nonlinear scaling. The recovery deficit *D*_rec_(*B*) (Eq. (9)) measures time-integrated loss of macroscopic coherence after task cessation. While *D*_rec_ increases overall with *B*, the scaling is weaker and more irregular (*β* ≈ 0.18, low *R*^2^), suggesting that integration loss does not grow proportionally with energetic cost.

**Figure 4.**
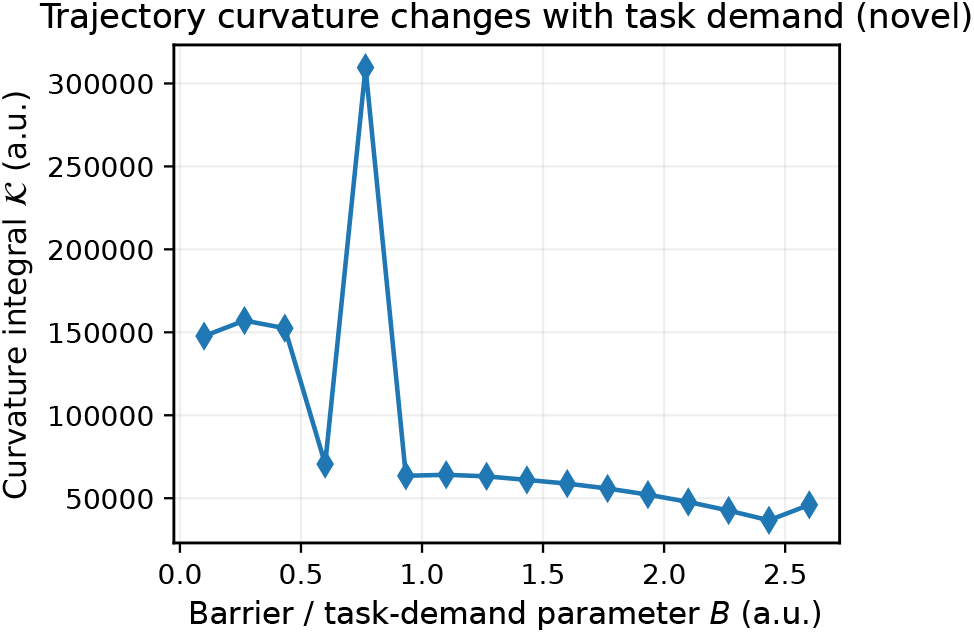
Mean-field trajectory curvature decreases with increasing demand. Integrated curvature 𝒦 (*B*) (Eq. (12)) quantifies geometric bending of the mean-field trajectory in the 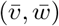 plane. Curvature exhibits negative scaling 𝒦 (*B*) ∼ *B*^−0.49^, indicating that higher-demand tasks produce more direct, less wandering collective transitions.

**Figure 5.**
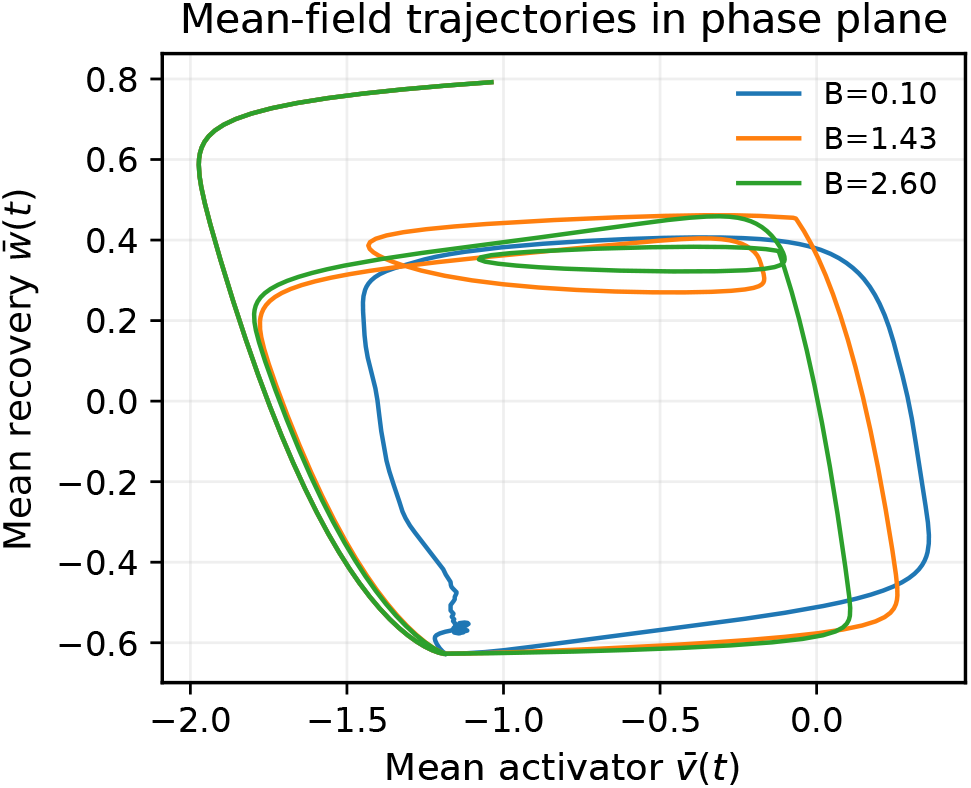
Representative mean-field trajectories for low, intermediate, and high demand. Trajectories in the 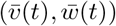 plane illustrate how increasing *B* alters collective state-space geometry. Low-demand regimes exhibit curved, exploratory excursions, whereas high-demand regimes produce more directed, ballistic transitions, consistent with curvature scaling.

**Figure 6.**
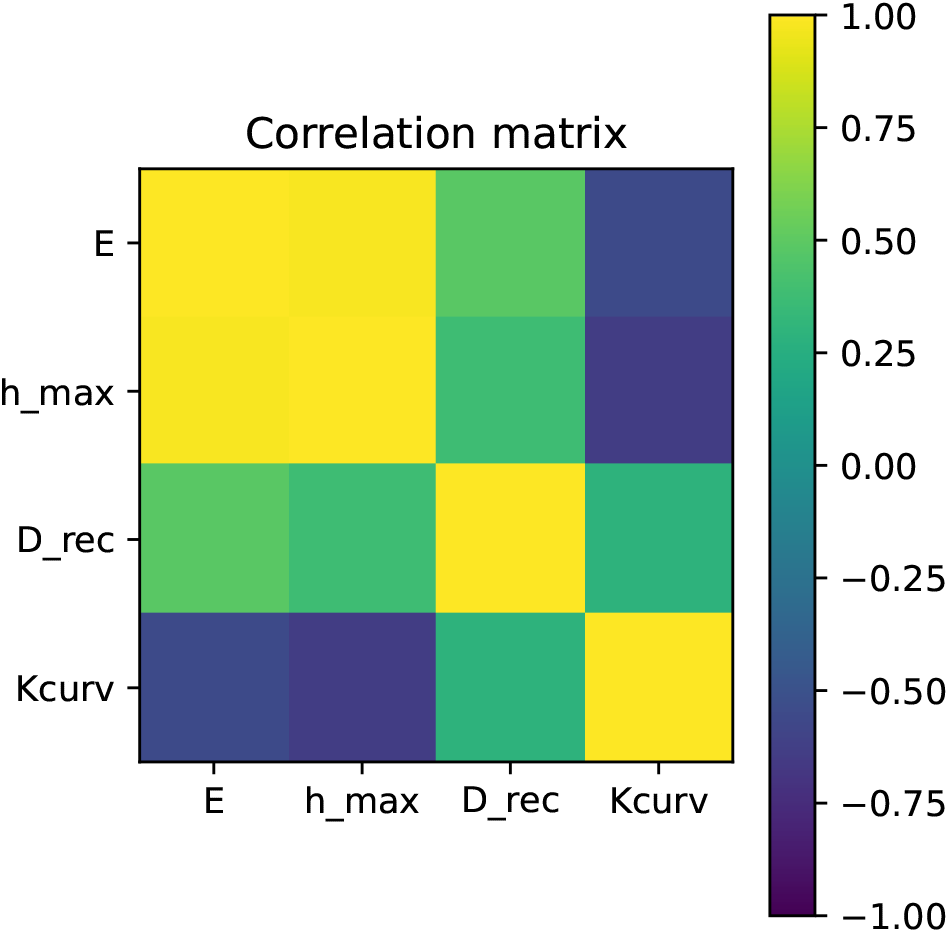
Correlation matrix of dynamical difficulty metrics. The full correlation structure among ε, *h*_max_, *D*_rec_, and 𝒦 reveals a dominant energetic–dispersion axis and partially independent recovery and geometric axes. Difficulty therefore occupies a structured region in a four-dimensional dynamical metric space rather than collapsing to a single scalar observable.

## 5 Discussion

### 5.1 Difficulty as dynamical reorganization

The present results suggest that task difficulty is not reducible to stimulus intensity or activation magnitude. Instead, difficulty emerges as structured reorganization of collective neural dynamics. Increasing task-demand parameter *B* induces coordinated changes across energetic, dispersive, integrative, and geometric dimensions of network activity.

The dominant axis identified here is the energetic–dispersion pair: transition action *E* and peak entropy *h*_max_ scale monotonically and remain strongly correlated. This indicates that harder tasks require greater dynamical work and recruit broader transient occupation of state space. In biophysical terms, difficulty corresponds to large-scale trajectory displacement across heterogeneous excitability landscapes.

Importantly, coherence recovery deficit *D*_rec_ does not scale proportionally with energetic cost. This partial independence suggests that integration loss reflects a distinct mechanism — namely, the transient fragmentation of modular coordination under conflict. Hard tasks therefore do not simply amplify activity; they reorganize coordination structure.

Finally, geometric curvature 𝒦 decreases with increasing demand, implying that low-demand regimes exhibit exploratory, curved trajectories, whereas high-demand regimes produce more directed state transitions. This geometric finding reveals that difficulty modifies the topology of collective trajectories, not merely their amplitude.

Taken together, these observations indicate that difficulty corresponds to a structured low-dimensional manifold embedded in the four-dimensional metric space

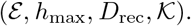

Thus, difficulty should be interpreted as a coordinated deformation of neural state-space geometry.

### 5.2. Biophysical interpretation

From a dynamical-systems perspective, increasing *B* effectively tilts the underlying vector field through modular conflict, shifting the network across competing attractor basins. Energetic scaling reflects increased traversal distance in state space. Entropy growth reflects heterogeneity amplification under forcing. Recovery deficit reflects the time required to restore collective phase coherence after perturbation. Curvature captures geometric redirection in the mean-field manifold.

This framework aligns with theories of cognitive effort as reconfiguration cost in distributed neural systems. However, the present formulation goes further by decomposing difficulty into independent dynamical components. In this sense, difficulty is not a scalar resource but a structured reorganization process.

### 5.3. Implications for artificial systems

Artificial neural networks typically measure difficulty through loss magnitude, gradient norm, or training time. These proxies implicitly capture energetic cost but neglect dispersion and integration structure. The present framework suggests that AI difficulty estimation could instead be grounded in dynamical metrics such as trajectory curvature, entropy of activation distributions, and recovery of internal synchronization following perturbations.

In particular, training steps that produce large dispersion without corresponding integration may signal structurally difficult transitions. Conversely, highly directed (low-curvature) transitions with large energetic cost may reflect constraint-driven optimization rather than exploratory difficulty. This multidimensional interpretation offers a potential bridge between biological computation and adaptive AI difficulty estimation.

### 5.4. Limitations and future directions

The present model employs minimal FitzHugh–Nagumo units and a simplified modular conflict structure. Real cortical networks exhibit higher-dimensional dynamics, plasticity, stochasticity, and hierarchical organization. Extending the framework to spiking models, adaptive synapses, or data-driven connectomes would allow direct empirical testing.

Furthermore, the subjective experience of difficulty was not modeled here. Instead, we focused on objective dynamical signatures. A key future step is to examine whether behavioral measures of difficulty correlate preferentially with energetic, dispersive, integrative, or geometric axes.

## 6 Conclusion

We have proposed a dynamical manifold theory of task difficulty grounded in heterogeneous excitable networks. By systematically varying a structured task-demand parameter, we demonstrated that difficulty does not collapse to a single scalar measure. Instead, it manifests as coordinated variation across energetic cost, state-space dispersion, coherence recovery, and geometric trajectory curvature.

These results suggest that cognitive difficulty corresponds to structured reconfiguration of neural state space. This perspective provides a mechanistic foundation for understanding effort, conflict, and problem hardness in biological systems and offers a principled pathway for difficulty estimation in artificial neural networks.

Difficulty, in this view, is not merely “more activation.” It is the geometry of reorganization.

## References

[1] Roger Ratcliff and Gail McKoon. The diffusion decision model: Theory and data for two-choice decision tasks. Neural Computation, 20(4):873–922, 2008.

[2] Joshua I Gold and Michael N Shadlen. Neural basis of decision making. Annual Review of Neuroscience, 30:535–574, 2007.

[3] Ian Goodfellow, Yoshua Bengio, and Aaron Courville. Deep Learning. MIT Press, 2016.

[4] Eugene M Izhikevich. Dynamical Systems in Neuroscience. MIT Press, 2007.

[5] Peter Dayan and L F Abbott. Theoretical Neuroscience. MIT Press, 2001.

[6] Michael Breakspear. Dynamic models of large-scale brain activity. Nature Neuroscience, 20:340–352, 2017.

[7] Gustavo Deco, Viktor Jirsa, and Anthony R McIntosh. Emerging concepts for the dynamical organization of resting-state activity. Nature Reviews Neuroscience, 12:43–56, 2011.

[8] David Attwell and Simon B Laughlin. An energy budget for signaling in the grey matter of the brain. Journal of Cerebral Blood Flow & Metabolism, 21:1133–1145, 2001.

[9] Karl Friston. The free-energy principle: a unified brain theory? Nature Reviews Neuroscience, 11:127–138, 2010.

[10] Yoshiki Kuramoto. Chemical Oscillations, Waves, and Turbulence. Springer, 1984.

[11] Juan A Acebrón, Luis L Bonilla, Conrad J Pérez Vicente, Felix Ritort, and Renato Spigler. The kuramoto model: A simple paradigm for synchronization phenomena. Reviews of Modern Physics, 77:137–185, 2005.

[12] Richard FitzHugh. Impulses and physiological states in theoretical models of nerve membrane. Biophysical Journal, 1:445–466, 1961.

[13] Jinichi Nagumo, Suguru Arimoto, and Shuji Yoshizawa. An active pulse transmission line simulating nerve axon. Proceedings of the IRE, 50:2061–2070, 1962.

[14] Lars Onsager and Stephen Machlup. Fluctuations and irreversible processes. Physical Review, 91(6):1505–1512, 1953.

[15] Mark I Freidlin and Alexander D Wentzell. Random Perturbations of Dynamical Systems. Springer, 3 edition, 2012.

[16] Claude E Shannon. A mathematical theory of communication. Bell System Technical Journal, 27(3):379–423, 1948.

[17] Thomas M Cover and Joy A Thomas. Elements of Information Theory. Wiley, 2 edition, 2006.

[18] Steven H Strogatz. From kuramoto to crawford: exploring the onset of synchronization in populations of coupled oscillators. Physica D: Nonlinear Phenomena, 143(1–4):1–20, 2000.

[19] William H Press, Saul A Teukolsky, William T Vetterling, and Brian P Flannery. Numerical Recipes: The Art of Scientific Computing. Cambridge University Press, 3 edition, 2007.

